# Resistance exercise protects mice from protein-induced fat accretion

**DOI:** 10.1101/2022.10.18.512689

**Authors:** Michaela E. Trautman, Leah N. Braucher, Christian Elliehausen, Wenyuan G. Zhu, Esther Zelenovskiy, Madelyn Green, Michelle M. Sonsalla, Chung-Yang Yeh, Troy A. Hornberger, Adam R. Konopka, Dudley W. Lamming

## Abstract

Low protein (LP) diets extend the lifespan of diverse species, and are associated with improved metabolic health in both rodents and humans. Paradoxically, many athletes and bodybuilders consume high protein (HP) diets and protein supplements, yet are both fit and metabolically healthy. Here, we examine this paradox using weight pulling, a validated progressive resistance exercise training regimen, in mice fed either an LP diet or an isocaloric HP diet. We find that despite having lower food consumption than the LP group, HP-fed mice gain significantly more fat mass than LP-fed mice when not exercising, while weight pulling protected HP-fed mice from this excess fat accretion. The HP diet augmented exercise-induced hypertrophy of the forearm flexor complex, and weight pulling ability increased more rapidly in the exercised HP-fed mice. Surprisingly, exercise did not protect from HP-induced changes in glycemic control. Our results confirm that HP diets can augment muscle hypertrophy and accelerate strength gain induced by resistance exercise without negative effects on fat mass, and also demonstrate that LP diets may be advantageous in the sedentary. Our results highlight the need to consider both dietary composition and activity, not simply calories, when taking a precision nutrition approach to health.

## Introduction

In contrast to the long-standing idea that “a calorie is a calorie,” research over the last two decades has shown that calories from different macronutrient sources have distinct impacts on health (Hall et al., 2015). Dietary protein in particular has been shown to have a critical role in the regulation of metabolic health and longevity; low protein (LP) diets are associated with reduced rates of diabetes and other age related diseases in human longitudinal studies, and promote leanness and insulin sensitivity in human randomized clinical trials of metabolically unhealthy adults (Ferraz-Bannitz et al., 2022; Fontana et al., 2016; Levine et al., 2014; Sluijs et al., 2010). In mice, low protein, high carbohydrate diets reduce adiposity, improve glucose tolerance, and extend lifespan (Hill et al., 2022; Richardson et al., 2021; Solon-Biet et al., 2014; Solon-Biet et al., 2015).

The idea that low protein diets are beneficial – and high protein diets are deleterious – challenges conventional wisdom. Several high-protein “fad” weight-loss diets have been popularized over the last two decades, and many researchers and medical professionals support consuming more protein to support healthy aging (Rodriguez, 2015). In addition to promoting satiety, some short-term studies have found that high protein intake improves glucose control in adults with type 2 diabetes (Dong et al., 2013; Gannon et al., 2003; Seino et al., 1983), and high protein diets are recommended for physically active individuals to support muscle growth and strength improvements (Andersen et al., 2005; Willoughby et al., 2007). Both exercise and amino acids activate the mechanistic target of TOR (mTOR) protein kinase, which stimulates the protein synthesis machinery needed to stimulate skeletal muscle hypertrophy (Schiaffino et al., 2021; Simcox and Lamming, 2022). To support this process, The Academy of Nutrition and Dietetics recommends consuming 1.2-2.0 grams of protein per kg of body weight (BW) per day in physically active individuals (Thomas et al., 2016). These protein estimations are substantially higher than the Recommended Daily Allowance (RDA) for sedentary people of 0.8 g/kg body weight, an amount intended to provide sufficient nutrients in 97.5% of people, but it remains controversial if protein intake above this level is detrimental in this population.

If high dietary protein intake was always deleterious, the many individuals consuming high protein diets or taking widely sold protein or BCAA supplements to maximize muscle growth following resistance training would have higher adiposity and an increased risk of diabetes. However, there is no data available to support such a surprising conclusion, and indeed the opposite is true: exercise reduces the risk of developing type 2 diabetes (Grontved et al., 2012). We are therefore left with a paradox: while the human and animal data suggesting that increased dietary protein intake is detrimental for metabolic health and increases the risk for numerous age-related diseases is robust, some of the people consuming the highest levels of protein – athletes – are metabolically very healthy.

Here, we examine the possibility that exercise can protect mice from the deleterious metabolic effects of a high protein diet normally observed in sedentary mice. In order to evaluate the metabolic effects of exercise and dietary protein intake independent of weight loss, we utilized a recently validated method of progressive resistance exercise for mice that does not cause weight loss (Zhu et al., 2021) to assess the interaction of dietary protein content and exercise in male C57BL/6J mice. We fed mice either a low protein (LP, 7% of calories from protein) or a high protein (HP, 36% of calories from protein) diet, and either pulled an increasing load of weight down a track 3x per week for 3 months, or pulled a sham unloaded cart. During the course of the experiment, we comprehensively assessed weight and body composition, metabolic parameters, and the fitness of the mice on these regimens. Finally, we euthanized the mice to determine the effect of both diet and exercise on tissue weight and muscle mass, muscle fiber type cross-sectional area and distribution, and muscle mitochondrial respiration.

In agreement with our hypothesis that HP diets impair metabolic health in sedentary mice, we found that HP-fed sham-exercised mice gained excess fat mass and increased in adiposity relative to LP-fed mice. In sharp contrast, HP-fed mice that engaged in resistance exercise showed hypertrophy of specific muscles and were protected from fat accretion. However, exercise did not protect HP-fed mice from the effects of protein on blood sugar control. Interestingly, while exercising HP-fed mice gained strength more rapidly than exercising LP-fed mice, the difference in maximum weight that could be pulled by each mouse was not significant, but both the HP diet and weight training regimen increased muscle diameter and in general, size. Interestingly, neither diet nor exercise altered mitochondrial respiratory capacity. Our research shows that resistance exercise protects from HP-induced increases in adiposity in mice, and suggests that metabolically unhealthy sedentary individuals consuming a HP diet or protein supplements might benefit from either reducing their protein intake or beginning a resistance exercise program.

## Results

### Resistance exercise protects against high-protein induced fat and body weight gain

Seven-week-old C57BL/6J mice were randomized to groups of equal weight and body composition, and placed on either a Low Protein (LP, 7% of calories from protein) or High Protein (HP, 36% of calories from protein) diet. Of note, while the protein content of rodent chow varies, typically 17-24% of calories are derived from protein (Tuck et al., 2020). The two diets were isocaloric, as calories from fat were kept at 19%, and carbohydrates were reduced in the HP diet to compensate for the increased calories from amino acids. We have previously utilized this same LP diet, and the full composition of both diets can be found in **Table 1**. Throughout the course of the study, weight was collected weekly and body composition was analyzed every three weeks via EchoMRI (**Fig. 1A**). Starting after five weeks on the diets, we exercised half of the mice on each diet using a recently validated mouse model of resistance exercise training, weight pulling (WP), in which mice pull progressively heavier weights down a track; the other half of the mice were sham-exercised by pulling an unloaded cart.

**Table 1.**
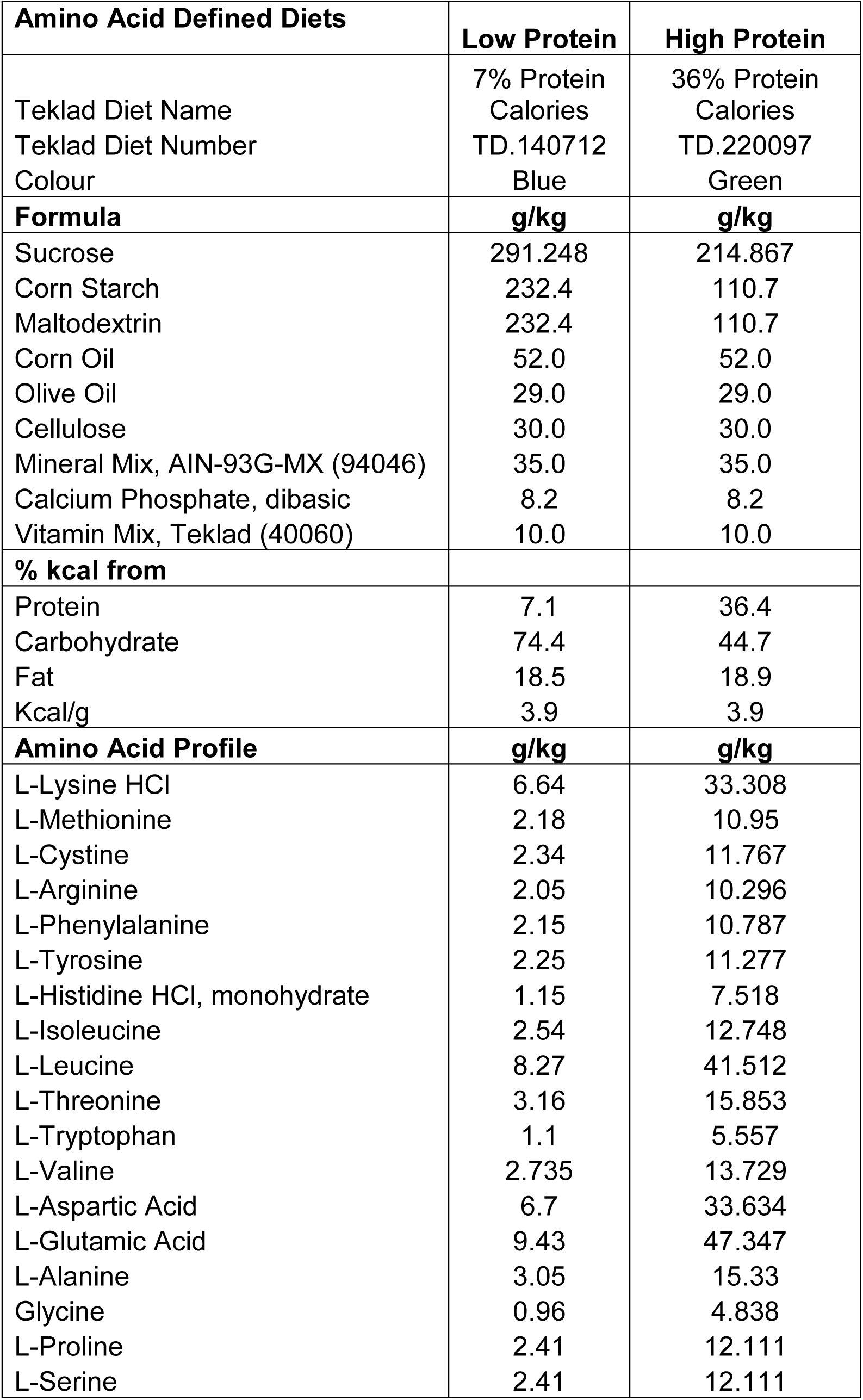
Diet composition.

**Figure 1.**
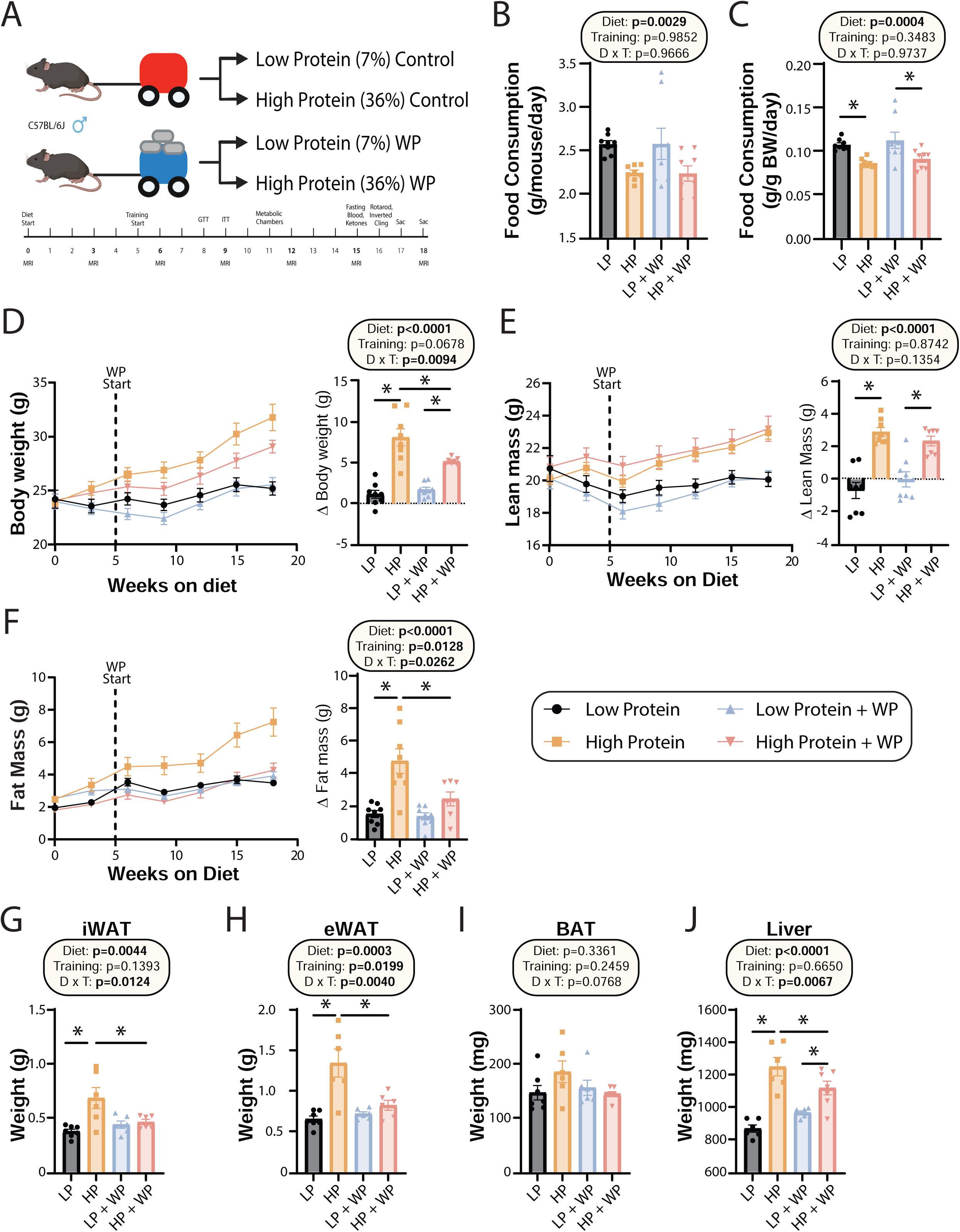
Weight pulling protects from high protein diet induced weight and fat gain. (A) Experimental design. (B-C) Food consumption per mouse (B) or normalized to body weight (C) after ∼6 weeks on the indicated diets. n=8/group. (D-F) Body weight (D), lean mass (E) and fat mass (F) over time, and change (Δ) from the beginning to end of study. n=7-8 mice/group. (G-J) Weight of the iWAT (G), eWAT (H), BAT (I), and liver (J) at the conclusion of the study. n=6-7 mice per group. (B-J) Statistics for the overall effects of diet, training, and the interaction represent the p value from a two-way ANOVA; *p<0.05, Sidak’s post-test examining the effect of parameters identified as significant in the 2-way ANOVA. Data represented as mean ± SEM.

On both a per-animal and a weight-normalized basis, LP-diet fed mice ate more than HP-fed mice, consistent with the satiating effect of dietary protein (**Fig. 1B-C**). There was no effect of WP on food consumption (**Fig. 1B-C**). Despite consuming fewer calories than LP-fed mice – the two diets have the same caloric density – HP-fed mice gained more weight over the course of 18 weeks. By the end of the experiment, HP-fed WP mice had gained significantly less weight than HP-fed sham-exercised controls (**Fig. 1D**). As we expected based on previous studies investigating the effects of a LP diet as well as data demonstrating the beneficial effects of protein on muscle growth (Andersen *et al*., 2005; Willoughby *et al*., 2007), HP-fed mice gained significantly more lean mass than LP-fed mice, although surprisingly exercise did not affect this gain (**Fig. 1E**). In accordance with our previous results, we found that sham-exercised HP-fed mice gained substantially more body fat than their LP-fed counterparts; however, we found that WP was protective against these effects (**Fig. 1F**).

At the conclusion of the experiment, we euthanized the animals and measured the weight of several fat depots and the liver. We found that the inguinal white adipose tissue (iWAT) and epididymal white adipose tissue (eWAT) were significantly heavier in sham-exercised HP-fed mice than in WP HP-fed mice (**Figs. 1G-H**). While there was no statistically significant effect on the mass of brown adipose tissue (BAT), the weight of this depot also trended higher in sham-exercised HP-fed mice than in WP HP-fed mice (**Fig. 1I**). Finally, there was a statistically significant effect of diet on liver weight, with HP-fed mice having heavier livers, an effect that was larger in sham-exercised mice than in WP mice (**Fig. 1J**).

We collected portions of the liver and iWAT for Oil RedO and H&E staining, respectively, and quantified the size and number of hepatic lipid droplets and adipocytes. Despite the increased liver weight of the HP groups, they did not have increased lipid accumulation; instead, there was a non-significant trend towards both larger and a greater number of liver lipid droplets in the LP-fed groups, with no effect of weight pulling (**Figs. 2A-C**). There was no difference in average iWAT adipocyte size or number (**Figs. 2D-F**).

**Figure 2.**
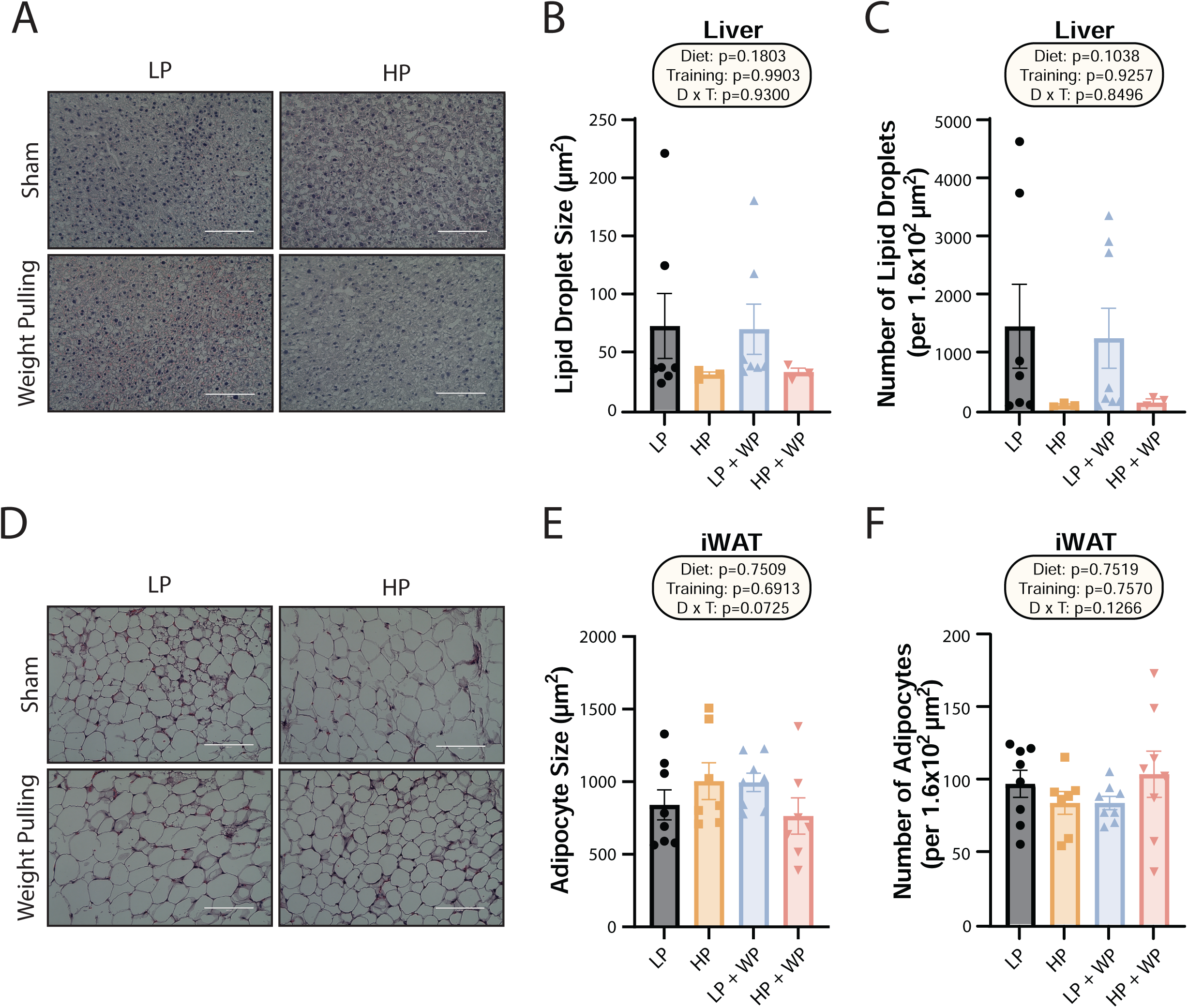
Dietary protein content and resistance training did not significantly impact liver lipid droplet or inguinal white adipocyte size. (A-C) Representative Oil-Red-O stained liver sections from mice in the indicated groups, with quantification of average lipid droplet size (B) and number (C). n=3-7/group. (D-F) Representative H&E stained iWAT sections from mice in the indicated groups, with quantification of average lipid droplet size (E) and number (F). n = 7-8/group. (B-C, E-F) Statistics for the overall effects of diet, training, and the interaction represent the p value from a two-way ANOVA; *p<0.05, Sidak’s post-test examining the effect of parameters identified as significant in the 2-way ANOVA. Scale bar = 100µM. Data represented as mean ± SEM.

### Mice fed a low protein diet have better glycemic control and higher energy expenditure than high protein-fed mice

We and others have previously shown that consumption of a low protein diet improves glucose tolerance and insulin sensitivity, while the literature suggests that resistance exercise improves insulin sensitivity in mice and humans (Cui et al., 2020; Fontana *et al*., 2016; Green et al., 2022; Kullmann et al., 2022; Laeger et al., 2014; McLeod et al., 2019; Westcott, 2012; Yu et al., 2021). We performed glucose and insulin tolerance tests; in agreement with our previous findings, we found that LP-fed mice had improved glucose tolerance relative to HP-fed mice, and there was a trend (p=0.0788) towards a positive effect of WP on glucose tolerance (**Fig. 3A**). Similarly, while LP-fed mice were more insulin sensitive than HP-fed mice under sham conditions, we did not observe a difference between LP and HP-fed mice that performed WP (**Fig. 3B**).

**Figure 3.**
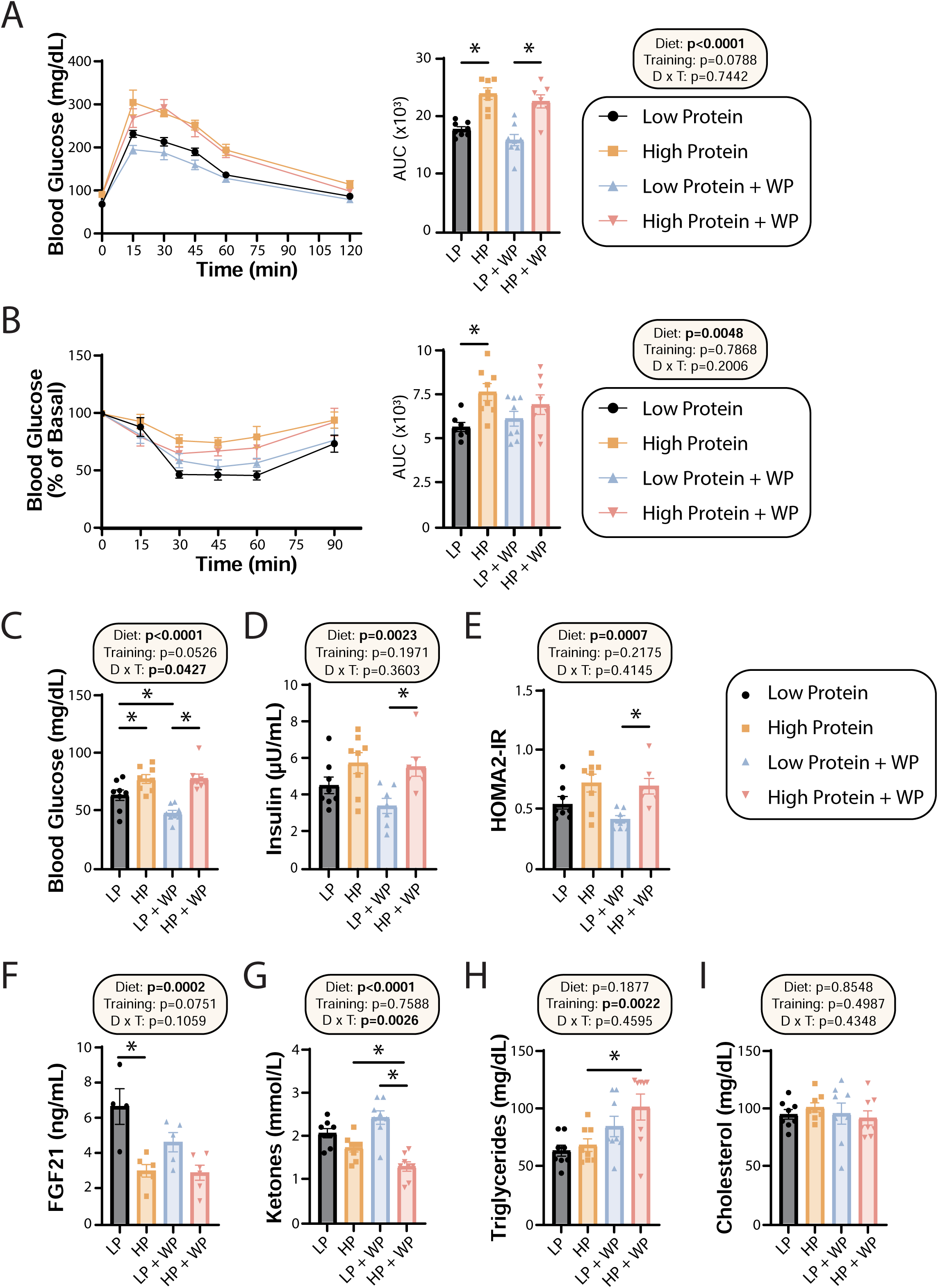
Effect of diet and exercise on glycemic control and blood metabolites. (A-B) Glucose (A) and Insulin (B) tolerance tests were performed after 9-10 weeks on the diet, respectively, and area under the curve (AUC) was calculated. n=7-8 mice/group. (C-E) Blood was collected from animals after a 16-hour overnight fast; fasting blood glucose (C) and insulin (D) were determined and used to calculate HOMA2-IR (E). n=7-8 mice/group. (F-I) Blood was collected from animals after a 16-hour overnight fast after 16 weeks on the diets. Fasting FGF21 (F), ketones (G), triglycerides (H) and cholesterol (I) were determined. n=4-8 mice/group. (A-I) statistics for the overall effects of diet, training and the interaction represent the p value from a two-way ANOVA, *p<0.05, Sidak’s post-test examining the effect of parameters identified as significant in the 2-way ANOVA. Data represented as mean ± SEM.

In agreement with a model in which high dietary protein is deleterious to sedentary animals, we observed that HP-fed mice had higher fasting blood glucose than LP-fed mice under sham exercise conditions (**Fig. 3C**). Interestingly, WP did not correct this difference – instead, we observed that there was an overall effect of HP-feeding towards increased fasting blood glucose level in both sham-exercised and WP mice (**Fig. 3C**). A similar effect of diet was noted on fasting insulin levels, with HP-fed WP mice having significantly higher insulin levels than LP-fed WP mice (**Fig. 3D**). In agreement with the overall effect of diet on fasting glucose and insulin levels, we also observed an overall effect of diet on insulin sensitivity calculated via the HOMA2-IR method, with LP WP mice having significantly better insulin sensitivity than HP-fed WP mice (**Fig. 3E**).

We and others have previously shown that a LP diet increases fasting FGF21 levels in C57BL/6J male mice (Green *et al*., 2022; Hill *et al*., 2022), and we observed that LP-fed sham mice had significantly higher levels of FGF21 in their blood compared to their HP-fed counterparts (**Fig 3F**). Interestingly, WP mice had a strong overall trend (p=0.0751) towards lower FGF21 levels, and LP-fed WP mice did not have significantly higher FGF21 levels than HP-fed WP mice (**Fig 3F**). FGF21 promotes ketogenesis, and in agreement with our FGF21 data we observed an overall effect of diet on fasting blood ketone levels, with LP-fed mice having higher ketone levels than HP-fed mice (**Fig. 3G**). Intriguingly, there was a significant interaction of diet and exercise, with WP lowering ketone levels in HP-fed mice.

Finally, we examined the effect of diet and exercise on blood lipids. Some studies have shown altered lipid levels in mice and humans fed a LP diet (Trevino-Villarreal et al., 2018), and we considered it likely that WP would promote a healthy lipid profile. Surprisingly, while diet had no effect on triglyceride or lipid levels, there was an overall increase of triglycerides in WP-fed mice, which was statistically significant in HP-fed mice (**Fig. 3H**). There was no effect of diet or exercise on blood levels of cholesterol (**Fig. 3I**).

In order to learn more about the effects of the HP diet on weight and adipose gain, we used metabolic chambers to evaluate multiple components of energy balance, including food consumption, spontaneous activity and energy expenditure. In agreement with our prior home-cage observations and the well-known satiating effect of dietary protein, we observed that HP-fed mice tended to consume less food than LP-fed mice, although this did not reach statistical significance. (**Supplemental Fig. 1A**). There was no overall effect of exercise on spontaneous activity, but in the light phase there was an overall effect of diet, with LP-fed mice moving more (**Supplemental Fig. 1B**).

We examined respiratory exchange ratio (RER) by calculating the ratio of O_2_ consumed and CO_2_ produced; an RER of close to 1.0 indicates that carbohydrates are being preferentially utilized for energy production, while a value near 0.7 indicates that lipids are the predominant energy source (Bruss et al., 2010; Yu et al., 2019). As we anticipated, there was an overall effect of diet on RER, with mice consuming the LP diet having a higher RER during the dark cycle, reaching statistical significance in the WP groups (**Supplemental Figs. 1C-D**). In agreement with previous studies, we observed a significant effect of the LP diet on energy expenditure, both on a per animal basis and when weight was considered as a covariate (**Supplemental Figs. 1E-F**). Resistance exercise did not have an overall effect on either RER or energy expenditure.

### Training while consuming a LP diet does not compromise maximum strength or coordination

Prior dogma suggests that in order to maximize strength and muscle hypertrophy from resistance exercise, consumption of a large quantity of high-quality protein is necessary (Phillips et al., 2005; Thomas *et al*., 2016). In agreement with this, we observed an significant overall effect of diet on the maximum load mice were able to pull during their training sessions as well as the corresponding area under the curve (AUC) (**Fig. 4A**), consistent with HP-fed mice being able to pull more weight. However, the difference was smaller than we had anticipated, and was statistically significant only during the third week of the training regimen; the maximum weight pulled by LP-fed and HP-fed mice appeared to converge after approximately 10 weeks of training (**Fig. 4A**). There was no overall effect of diet on the average number of sets per training bout (**Fig. 4B**) or the average number of stimulatory touches by the investigator (**Fig. 4C**).

**Figure 4.**
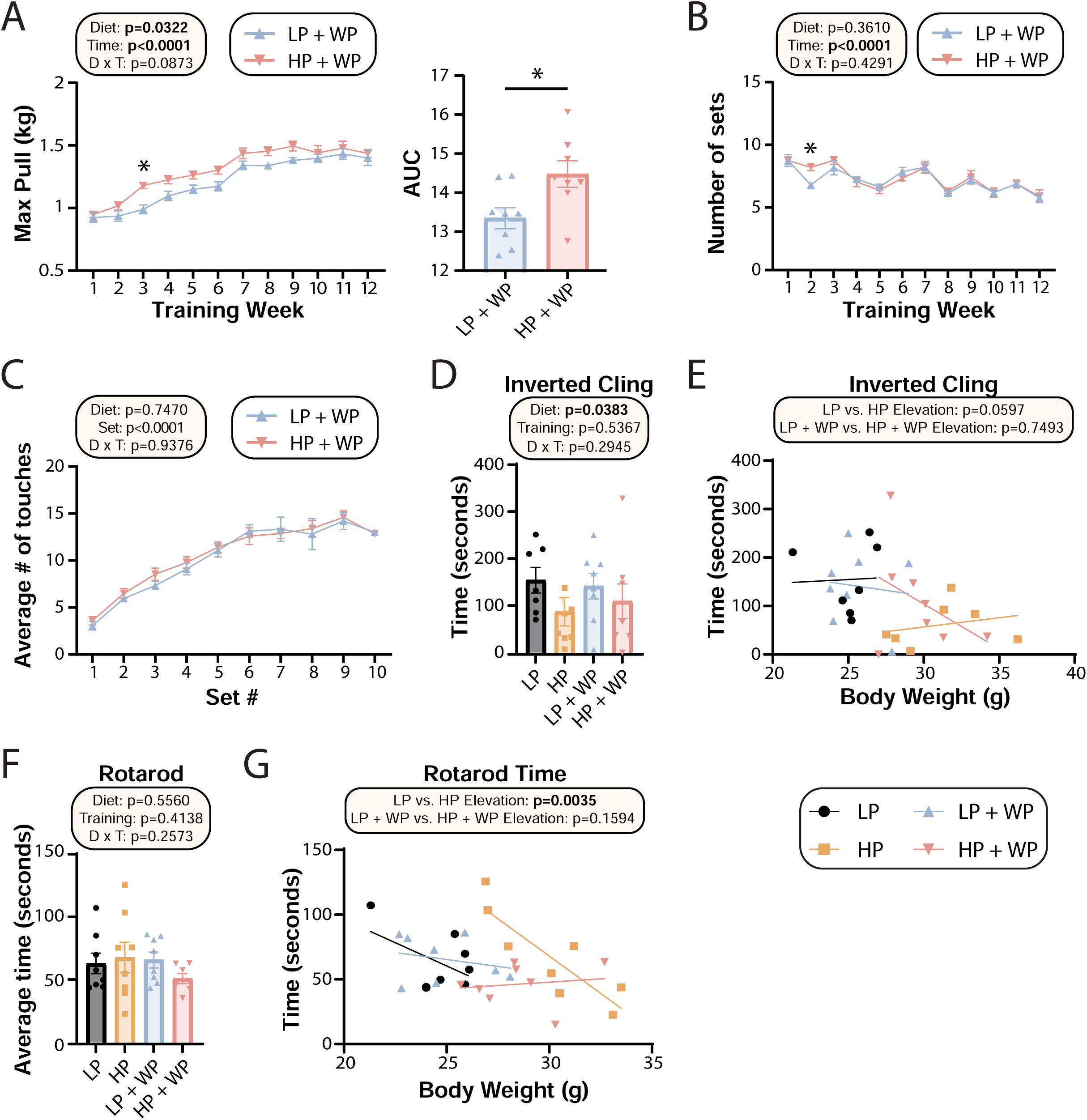
Strength and muscle growth is maximized by high protein and progressive resistance exercise. (A-C) Weight pulling was performed three times per week for 12 weeks. The average maximum weight pulled each week with area under the curve (AUC) (A), and the number of sets achieved (B) is shown. Similarly, the number of stimulatory touches (C) by the investigator on all mice was averaged through multiple weeks of training and plotted per set. (A-C) n=8 mice/group (A-B); for (C) values for 8 mice/group were assayed over n=5 weeks. Statistics for the overall effects of diet, time or set, and the interaction represent the p value from a two-way RM ANOVA or REML; *p<0.05 from a Sidak’s post-test examining the effect of parameters identified as significant in the 2-way ANOVA. (D-G) Physical performance was assessed by an inverted cling test (D-E) or rotarod test (F-G). (E, G) Cling time (E) or rotarod time (G) as a function of body weight was calculated (data for each individual mouse are plotted, and slopes and intercepts were calculated using ANCOVA). n=7-8 mice/group. Statistics for the overall effects of diet, training and the interaction represent the p value from a two-way ANOVA, *p<0.05, Sidak’s post-test examining the effect of parameters identified as significant in the 2-way ANOVA. Data represented as mean ± SEM.

The strength gains from WP did not translate into improved performance on two other tasks assessing muscle strength and function, inverted cling and rotarod (**Fig. 4D-G**). We observed an overall positive effect of a LP diet on inverted cling time, an effect that persisted (p=0.0597) in sham-exercised mice when we analyzed inverted cling performance with weight as a covariate (**Fig. 4D**). Although we observed no overall effect of either diet or training on raw rotarod performance (**Fig. 4F**), we observed a significant positive effect of a HP diet on the rotarod performance of sham-exercised mice when weight was considered as a covariate (**Figs. 4G**).

### Weight pulling and high protein diet promote muscle hypertrophy without impacting mitochondrial respiration

At the conclusion of the weight training regimen and the *in vivo* experiments described above, we euthanized the animals and collected numerous tissues. As noted above, WP in HP-fed mice reduced the mass of iWAT, eWAT, and liver (**Figs. 1F-I**). We also collected a diverse array of skeletal muscles from different regions of the body. We have previously shown that weight pulling induces hypertrophy in the flexor digitorum longus (FDL) (Zhu *et al*., 2021), and as we expected we found an overall positive effect of both a HP diet and weight pulling on FDL mass, both in absolute terms and when normalized to either body mass or tibia length (**Figs. 5A-C**). We observed the greatest muscle mass in mice that completed the WP regimen while consuming a HP diet, but HP diet consumption alone also had a positive effect on absolute FDL mass and FDL mass normalized to tibia length (**Figs. 5A-C**).

**Figure 5.**
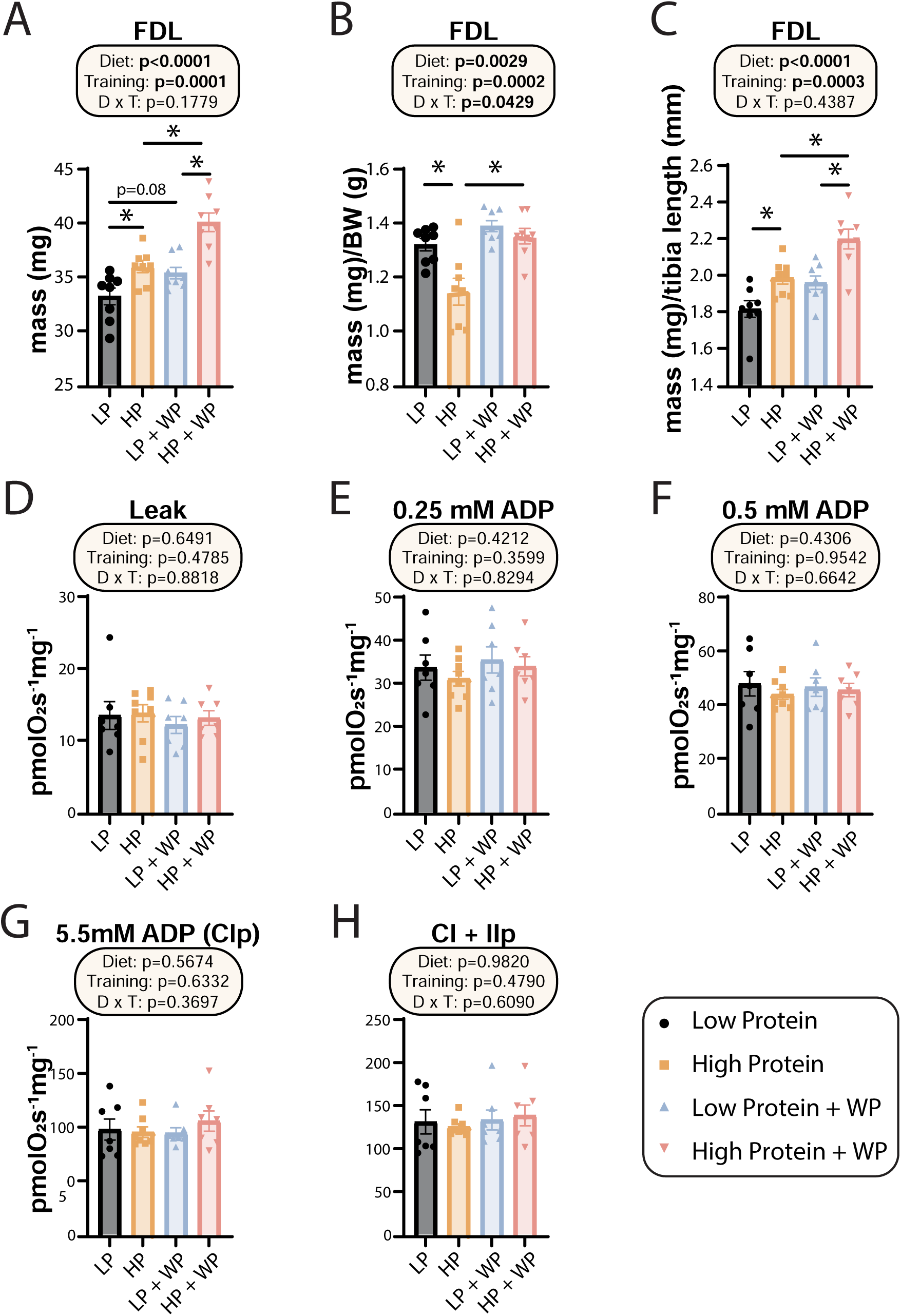
Weight pulling and high protein diet increased FDL mass but not mitochondrial respiration. (A-C) The muscle mass of the FDL in absolute mass (A), normalized to body weight (B), and (C) normalized to tibia length. n=8/group. (D-H) Mitochondrial respiration parameters as measured in the FDL, including mitochondrial leak (D), and following addition of 0.25 mM (E), 0.5 mM (F), and 5.5 mM ADP (G). (H) 10mM succinate was provided to evaluate complex I and II driven respiration. n=7-8 mice/group. (A-H) Statistics for the overall effects of diet, training and the interaction represent the p value from a two-way ANOVA, *p<0.05, Sidak’s post-test examining the effect of parameters identified as significant in the 2-way ANOVA. Data represented as mean ± SEM.

In humans, mitochondrial respiration is increased after both aerobic and resistance training regimens (Konopka et al., 2015; Konopka et al., 2019; McKenna et al., 2022; Robinson et al., 2017), perhaps due to the high energetic demands imposed by protein synthesis (Rolfe and Brown, 1997; Waterlow, 1984), which is upregulated following resistance training (Ogasawara et al., 2016). However, the mitochondrial response to varying amounts of dietary protein with or without resistance training has not been examined, and we therefore performed high-resolution respirometry in permeabilized muscle fibers from the FDL. Using complex-I linked substrates pyruvate, glutamate, and malate we found no change to leak respiration (**Fig. 5D**). We next evaluated sub saturating and saturating complex-I driven respiration by providing ADP at 0.25mM, 0.5mM and 5.5mM; none of these parameters were different in any intervention group (**Figs. 5E-G**). Furthermore, we saw no difference in any group following succinate addition for complex I+II linked respiration (**Fig. 5H**).

We examined the effect of diet and weight pulling on the mass of the quadriceps, soleus, plantaris, and the forearm flexor complex, measuring mass in absolute terms (**Supplemental Figs. 2A-D**) and normalized to tibia length (**Supplemental Figs. 2E-H**). Overall, there was no positive effect of either resistance exercise or high protein diet on quadricep mass (**Supplemental Figs. 2A, E**), while the mass of the soleus and plantaris muscles were positively impacted by dietary protein but largely resistant to the benefits of WP (**Supplemental Figs. 2B-C, F-G**). However, there were strong effects of WP and a significant interaction between dietary protein and exercise on the mass of the forearm flexor complex, with the HP-fed WP mice having the greatest forearm flexor complex mass in raw weight and when normalized to tibia length (**Supplemental Figs. 2D, H**).

Finally, we assessed the maximum diameter of the bicep and forearm (**Fig. 6A**) and found that there were clear overall effects of both dietary protein and weight pulling on bicep and forearm diameter (**Fig. 6B-C**). Bicep diameter was maximized under high protein weight pulling conditions (**Fig. 6A-B**). There was a less potent but still significant effect on the forearm; HP-fed WP mice had the largest forearm diameter of all groups (**Fig. 6C**). Conversely, non-exercising LP-fed mice had the smallest bicep and forearm diameter (**Figs. 6A-C**).

**Figure 6.**
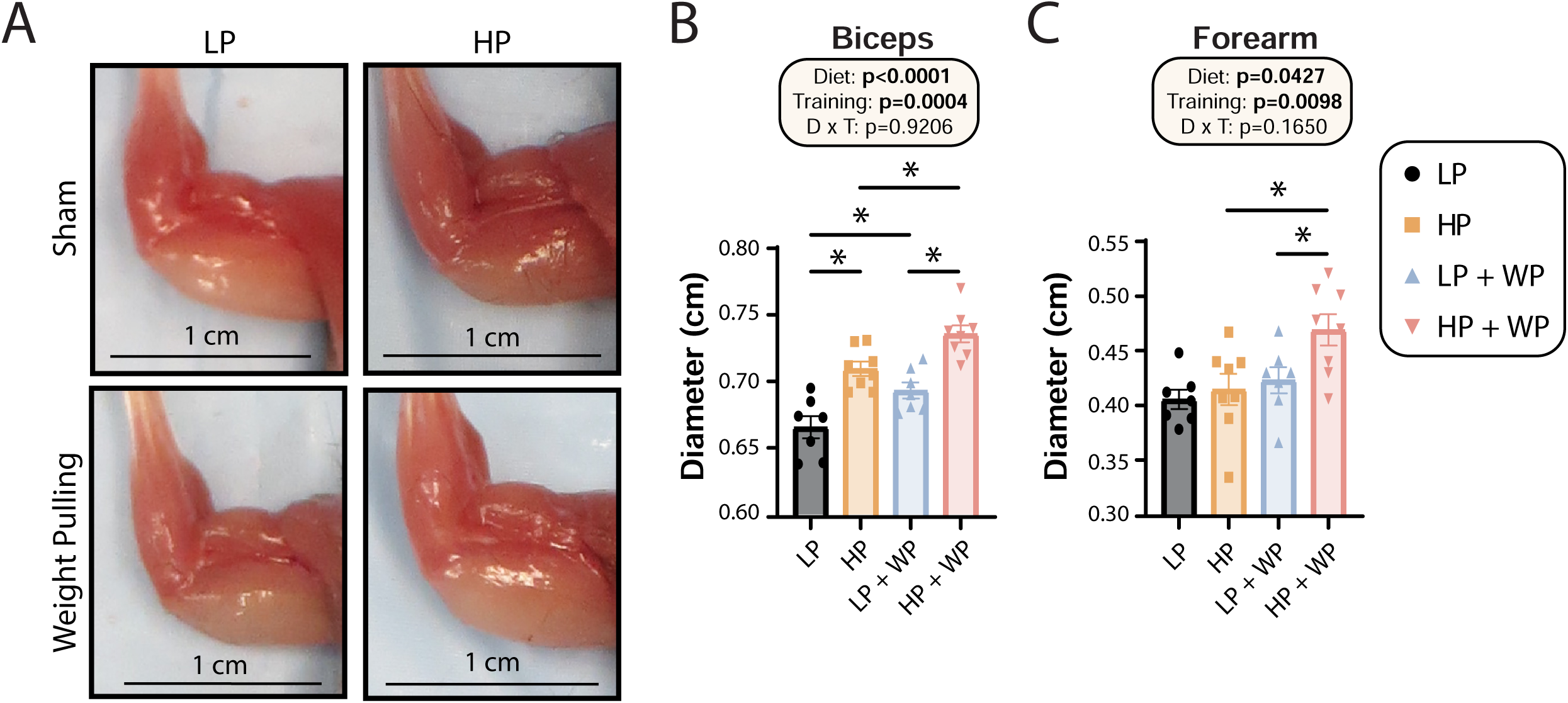
Bicep and forearm hypertrophy is maximized by HP diets and WP. (A-C) Representative images of arm musculature (A) with quantification of biceps (B) and forearm (C) diameter. (B-C) n=7-8 mice per group. Statistics for the overall effects of diet, training and the interaction represent the p value from a two-way ANOVA, *p<0.05, Sidak’s post-test examining the effect of parameters identified as significant in the 2-way ANOVA. Data represented as mean ± SEM.

Next, we further analyzed the FDL through staining (**Fig. 7A**) and quantification (**Fig. 7B-H**). There was no significant difference in mid-belly cross-sectional area (CSA) or the number of fibers per cross section (**Fig. 7B-C**), indicating that alterations in dietary protein or weight pulling might not induce hyperplasia of muscle fibers (Jorgenson and Hornberger, 2019). There was, however, an overall effect of diet on the CSA of muscle fibers, with individual fibers in HP-fed mice having greater CSA (**Fig. 7D**).

**Figure 7.**
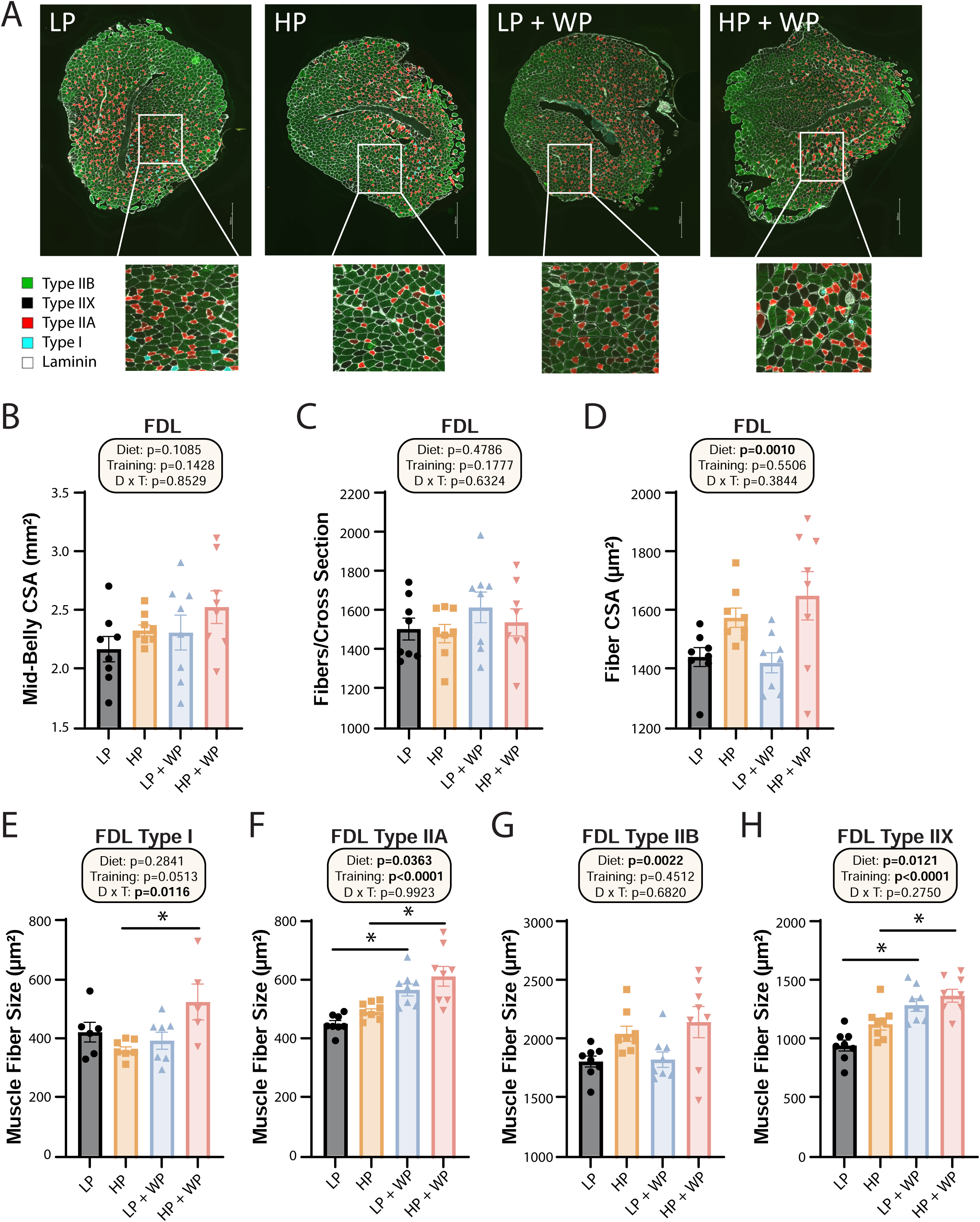
FDL fiber type hypertrophy is maximized by high protein diets and weight pulling. (A-H) Representative images of the FDL and fiber type with quantification of mid-belly CSA (B), fibers per cross section (C), fiber CSA (D), and individual muscle fiber type size: type I (E), type IIA (F), type IIB (G) and type IIX (H).n=5-8 mice per group. Statistics for the overall effects of diet, training and the interaction represent the p value from a two-way ANOVA, *p<0.05, Sidak’s post-test examining the effect of parameters identified as significant in the 2-way ANOVA. Data represented as mean ± SEM.

When we analyzed fiber type specific CSA, we observed an overall effect of diet on the size of type IIA, type IIB, and type IIX fibers, with HP diets increasing fiber size (**Figs. 7F-H**). We also saw an effect of resistance exercise on the size of type IIA and type IIX fibers, with WP inducing fiber hypertrophy (**Figs. 7F, 7H**). Finally, we observed a significant interaction between dietary protein and WP on the size of type I fibers, with the greatest hypertrophy occurring in HP-fed WP mice (**Figs. 7E**).

## Discussion

Dietary protein is a critical regulator of a wide variety of biological processes that determine metabolic health and lifespan in diverse species (Trautman et al., 2022). Consumption of protein is generally thought of as good, promoting muscle growth and strength particularly in combination with exercise, and skeletal muscle function and mass is associated with a host of health benefits including a reduced risk for diabetes and frailty (Liao et al., 2019; Phillips, 2007). Paradoxically though, in sedentary humans increased consumption of dietary protein is associated with cancer, cardiovascular disease, and diabetes as well as increased mortality (Lagiou et al., 2007; Levine *et al*., 2014; Mittendorfer et al., 2020; Sluijs *et al*., 2010; Zhang et al., 2020). In agreement with the potential for dietary protein to negatively impact health, low protein diets and diets with reduced levels of specific essential amino acids promote healthspan and lifespan in flies and rodents (Flores et al., 2022; Green et al., 2023; Juricic et al., 2020; Lee et al., 2014; Lees et al., 2014; Orentreich et al., 1993; Richardson *et al*., 2021; Solon-Biet *et al*., 2014), and short-term protein restriction improves the metabolic health of metabolically unhealthy adult humans (Ferraz-Bannitz *et al*., 2022; Fontana *et al*., 2016).

We know that many of the humans deliberately consuming high protein diets or consuming protein supplements to support their exercise regimen are not metabolically unhealthy – indeed, many of these individuals have commendable metabolic health (Antonio et al., 2015; Antonio et al., 2016; Grontved *et al*., 2012). We considered that a potential solution to this paradox is that exercise itself is protective from the effects of high protein diets; our and others previous studies examined sedentary mice and largely sedentary humans. Here, we investigated this hypothesis by feeding isocaloric diets with either high or low levels of dietary protein to mice and subjecting them to a recently validated mouse model of progressive resistance exercise, weight pulling (WP) or sham control exercise. Importantly, unlike aerobic exercise models like treadmill running, WP does not result in weight loss, allowing us to study the metabolic impact of exercise without confounding changes in body weight. We also took advantage of this opportunity to rigorously test long-standing assumptions regarding the effects of dietary protein and resistance exercise on muscle mass and function.

In agreement with previous studies by our lab and others, we found that in sham-exercised mice, LP-fed mice were metabolically healthier than HP-fed animals, remaining leaner and having better glycemic control; conversely, HP-fed animals accumulated fat mass and had worse glycemic control. However, when HP-fed mice were subjected to WP, the negative effects of the HP diet on overall fat mass gain as well as on individual fat depots were completely blocked. Surprisingly, although WP did further lower fasting blood glucose in LP-fed mice and these mice had a tendency towards improved glucose tolerance and lower fasting insulin, the negative effect of the HP diet on glucose tolerance and insulin sensitivity was not blocked by WP.

While HP-feeding did, as we expected, support muscle mass gain and strength gain more than an LP diet, the superiority of the HP diet in the maximum weight pulled group was transitory; by the end of the training period, LP-fed mice could pull just as much weight as HP-fed mice. There was a modest effect of diet on performance of mice on inverted cling and rotarod tasks, with HP-fed sham mice performing better on the rotarod than LP-fed sham mice when weight was considered as a covariate, but this effect was not observed in WP groups. As rotarod and inverted cling outcomes can be improved by voluntary aerobic exercise (wheel running) (Graber et al., 2015), this could be the result of either the type of exercise or the precise muscles strengthened by WP vs. aerobic exercise. Somewhat surprisingly, mitochondrial respiration outcomes were also independent of diet and exercise in the FDL, but we did not examine mitochondrial function in other tissues.

Our study was subject to a number of limitations. We examined only young inbred C57BL/6J male mice, and we have previously shown that sex, genetic background, and age impact the metabolic response to dietary protein (Green *et al*., 2022). We consider it likely that these factors may also influence the response to exercise, and examining the interaction between diet and WP in females, older mice, and different genetic backgrounds should be a high priority for future studies. Further, as the metabolic response to dietary protein is responsive to the exact level of protein in the diet, examining additional levels of dietary protein between the LP and HP diets we examined here might provide greater insight. This would also allow the study of dietary protein levels within the Acceptable Macronutrient Distribution Range (AMDR) for humans, which is 10-35% of calories from protein. Finally, to keep our diets isocaloric, isonitrogenous and consistent in fat content, we had to reduce carbohydrates in the HP diet, which might play a role in the results found here.

Many people who eat high protein diets or take protein supplements also engage in aerobic exercise, which we did not examine in the present study. Here, we have not examined the effects of dietary protein and exercise on molecular changes and have not explained how WP inhibits HP-induced gains in adiposity. The hormone FGF21, which is induced by a LP diet, is responsive to exercise in mice and humans (Haghighi et al., 2022; Jin et al., 2022; Li et al., 2022), but we did not observe increased FGF21 in the LP weight trained group, suggesting another mechanism could be at play. As for the improved body composition of HP-fed WP mice, we hypothesize that these interventions may activate beiging in iWAT or BAT. Other possibilities include changes in mitochondrial respiration in muscles other than the FDL, or exercise-induced alterations in lipogenesis or lipolysis. Lastly, we have not explored the role of individual amino acids in the response; many athletes and body builders take supplements enriched in BCAAs, and future work should determine if there is a specific role for these or other dietary amino acids in the metabolic and physiological response to resistance exercise.

Finally, when broken down to individual muscle fiber types, we saw an increase in type 1 muscle fiber size in the FDL of the HP + WP group, which indicates that both high protein and resistance training are necessary for inducing hypertrophy of slow-twitch, oxidative fiber types. However, as three of our HP + WP samples had no detectable type 1 fibers, future research should evaluate this effect in muscles with an abundance of type 1 fibers. These findings are consistent with specific fiber type hypertrophy previously observed in the FDL in response to WP (Zhu *et al*., 2021). We saw a similar induction of growth by both diet and training in type IIA fibers, which are fast-twitch, oxidative and glycolytic. Here, HP feeding resulted in larger fibers than LP in the context of both sham training and weight pulling, which is consistent with human recommendations that increased dietary protein supports muscle hypertrophy (Thomas *et al*., 2016). Surprisingly, there was no effect of training observed in fast-twitch, glycolytic type IIB fibers, but there was an effect of diet, with HP feedings inducing hypertrophy. Finally, fast-twitch type IIX fibers demonstrated a similar trend as type IIA fibers, with both a HP diet and WP inducing growth. A striking takeaway observed here is that there was no difference in hypertrophy of Type IIX and IIA fibers in the FDL induced by WP when fed a low or high protein diet. Though these changes occurred in just one muscle, and effects on other muscles may vary, our results indicate that both resistance training and high protein diets individually and in combination optimize growth of muscle fibers.

In summary, dietary protein and progressive resistance exercise interact to regulate metabolic health as well as strength and muscle mass. Specifically increasing protein consumption has – as we anticipated based on both animal and human data – negative effects on the metabolic health of mice that are not exercising. However, the negative effects of a HP diet on body composition are eliminated in mice performing progressive resistance exercise training. While mice consuming a HP diet gained strength during weight pulling more quickly than LP-fed mice, the ultimate max load the HP and LP-fed mice could pull by the end of twelve weeks was indistinguishable, despite clear differences in muscle mass. Future research could investigate this apparent paradox by examining the effect of the diet and WP regimens on muscle mass and fiber type at earlier time points than those examined in this study. However, our results are consistent with studies in healthy, resistance-trained individuals that show high protein diets can be consumed without negative effects on body composition (Antonio *et al*., 2015; Antonio *et al*., 2016). We hypothesize that further studies will find that sedentary individuals would be metabolically healthier if consuming a low protein diet. Finally, our results suggest that a precision nutrition approach to metabolic health must consider not only diet, sex, and genetic variation, but also take into account activity level.

## Methods

### Animal care, housing and diet

All procedures were performed in conformance with institutional guidelines and were approved by the Institutional Animal Care and Use Committee of the William S. Middleton Memorial Veterans Hospital. Male C57BL/6J mice were procured from the Jackson Laboratory (000664) at six weeks of age. All studies were performed on animals or on tissues collected from animals. All mice were acclimated to the animal research facility for at least one week before entering studies. All animals were housed 2 per cage in static microisolator cages in a specific pathogen-free mouse facility with a 12:12 h light–dark cycle, maintained at approximately 22 °C. At the start of the experiment, mice were randomized to receive either the 7% (Low Protein, TD.140712) or 36% (High Protein, TD.220097) amino acid defined diets, and assigned to the sham or weight pulling groups; all diets were obtained from Envigo. In the High Protein diet, carbohydrates were reduced and amino acids increased relative to the low protein diet in order to keep the diets isocaloric, while calories from fat were held fixed. Full diet descriptions, compositions and item numbers are provided in **Table 1**.

### Weight training paradigm

The procedures of this weight pulling regimen (acclimation, weighted paradigm, and unweighted paradigm) were described previously (Zhu *et al*., 2021).

### Metabolic Phenotyping

Glucose and insulin tolerance tests were performed following a 16 hour overnight or 4 hour fast, respectively and then injecting either glucose (1g/kg) or insulin (0.75U/kg) intraperitoneally (Bellantuono et al., 2020; Yu *et al*., 2019). Glucose measurements were taken using a Bayer Contour blood glucose meter and test strips. Blood ketone measurements were taken using a KetoBM ketone meter and test strips that detect beta-hydroxybutyrate, the most abundant ketone in the body and proxy for ketone level. Mouse body composition was determined using an EchoMRI Body Composition Analyzer. For assays of multiple metabolic parameters (O_2_, CO_2_, food consumption, and activity tracking), mice were acclimatized to housing in a Columbus Instruments Oxymax/CLAMS-HC metabolic chamber system for approximately 24 hours, and data from a continuous 24-hour period was then recorded and analyzed.

### Tissue collection

Terminal collections were performed 60–96 h after the final training session and after a 3 hour morning fast. During this procedure, all potentially identifiable information (e.g., tail markings, etc.) was masked, and then the animals were weighed and subsequently anesthetized with isoflurane. The plantaris, flexor digitorum longus (FDL), forearm flexor complex (FF) (which consisted of the flexor carpi ulnaris, flexor carpi radialis, flexor digitorum superficialis, and all three heads of the flexor digitorum profundus), and soleus (SOL) muscles from both the left and right hindlimb were then weighed. The left and right plantaris and right FDL were submerged in optimal cutting temperature compound (OCT, Tissue-Tek; Sakura Finetek, The Netherlands) at resting length, and frozen in liquid nitrogen-chilled isopentane. Left FDL was prepared for mitochondrial analysis. At this point, the mice were euthanized by cervical dislocation, a photograph that included both a scale bar and the musculature of the left forelimb was obtained, and then quadriceps muscles from both the left and right side of the body were collected. In addition to skeletal muscles, the inguinal white adipose tissue (iWAT), brown adipose tissue (BAT), epididymal white adipose tissue (eWAT), liver, as well as the right tibia bone, were collected post-mortem and frozen in liquid nitrogen or prepared for histology (OCT or formalin, to then be incubated in ethanol). Oil Red O staining of liver samples and H&E staining of iWAT samples were conducted by UW Carbone Cancer Center Experimental Animal Pathology Lab (UWCCC EAPL) on a fee-for-service basis. Muscle photographs were taken with a ruler for scale, and precise maximum muscle diameter was measured using ImageJ. All of the collection procedures were performed by investigators blinded to treatment and diet group.

### Immunohistochemistry and image analysis

Fiber type staining and analysis was performed as previously described (Zhu *et al*., 2021). ImageJ software was utilized to quantify hepatic lipid droplets and the ImageJ Adiposoft plugin was used to quantify iWAT adipocytes.

### High Resolution Respirometry

FDL muscle was excised, weighed, and cut in half. The distal half was snap frozen in liquid nitrogen while proximal half was placed in ice cold (4°C) Buffer X containing (in mM) 7.23 K2EGTA, 2.77 CaK2EGTA, 20 imidazole, 20 taurine, 5.7 ATP, 14.3 phosphocreatine, 6.56 MgCl2·6H2O, and 50 K-MES (pH 7.1). Muscle was mechanically permeabilized with fine tip forceps removing connective and adipose tissue. Fiber bundles were chemically permeabilized in Buffer X with saponin (50 μg/mL) for 30 min on ice. Permeabilized muscle fiber bundles were rinsed with MiR05 containing (in mM) 0.5 mM EGTA, 3 MgCl2, 60 K-lactobionate, 20 taurine, 10 KH2PO4, 20 HEPES, 110 sucrose, 1 g/L BSA essentially fatty acid free, (pH 7.1) and 25 μM blebbistatin. Fibers were then blotted on filter paper to remove excess buffer and weighed on a microscale. Fibers bundles (1.5-2.5 mg) were then placed into chambers of the Oxygraph-2k (O2K; OROBOROS INSTRUMENTS, Innsbruck, Austria) containing MiR05 plus 12.5 μM blebbistatin, a myosin ATPase inhibitor, at 37°C. Chambers were hyperoxygenated to ∼425uM and mitochondrial respiration was supported by complex-I linked substrates 5mM pyruvate, 10mM glutamate, and 0.5mM malate. ADP was provided at 0.25mM, 0.5mM and 5.5mM to evaluate sub saturating and saturating complex-I driven mitochondrial respiration. Mitochondrial membrane integrity was evaluated using 10μM cytochrome c. A >15% change in oxygen consumption was considered a loss of mitochondrial membrane integrity and 3 samples were removed from final analysis. 10mM succinate was provided to evaluate complex I and II driven respiration. Data is normalized to muscle fiber bundle wet weight.

### Statistics

Data are presented as the mean ± SEM unless otherwise specified. Statistical analyses were performed using one-way, two-way ANOVA, two-way RM ANOVA, or a mixed-effects model (Restricted Maximum Likelihood (REML)) followed by an appropriate posttest as specified in the figure legends. In all figures, n represents the number of biologically independent animals, except in figure 5C, where n represents the number of weeks that touches were determined for across all animals. Outliers were determined using GraphPad Prism Grubbs’ calculator (https://www.graphpad.com/quickcalcs/grubbs1/). Sample sizes were chosen based on our previously published experimental results with the effects of dietary interventions plus consideration of the training time required by personnel (Cummings et al., 2018; Fontana *et al*., 2016; Yu *et al*., 2021; Yu et al., 2018). Data distribution was assumed to be normal, but this was not formally tested.

## ACKNOWLEDGEMENTS

We thank Santosh Kumari for assistance with tissue collection. The Lamming Laboratory is supported in part by the NIH/National Institute on Aging (AG056771, AG062328, AG061635, AG081482 and AG084156), NIH/NIDDK (DK125859) and startup funds from the University of Wisconsin-Madison School of Medicine and Public Health and Department of Medicine to D.W.L. The Konopka lab is supported in part by the NIH/NIA (AG067464, AG076941, and AG081482). The Hornberger lab is supported in part by the NIH/NIAMS (AR057347 and AR074932). CY.-Y. is supported by a NIH/NIA F32 postdoctoral fellowship (AG077916). M.M.S. is supported by a Research Supplement to Promote Diversity in Health-Related Research (3RF1AG056771-06S1). The UWCCC Experimental Pathology Laboratory is supported by the University of Wisconsin Carbone Cancer Center Support Grant P30 CA014520. The Lamming laboratory was supported in part by the U.S. Department of Veterans Affairs (I01-BX004031 and IS1-BX005524), and this work was supported using facilities and resources from the William S. Middleton Memorial Veterans Hospital. The content is solely the responsibility of the authors and does not necessarily represent the official views of the NIH. This work does not represent the views of the Department of Veterans Affairs or the United States Government.

---

**Supplemental Figure 1.**
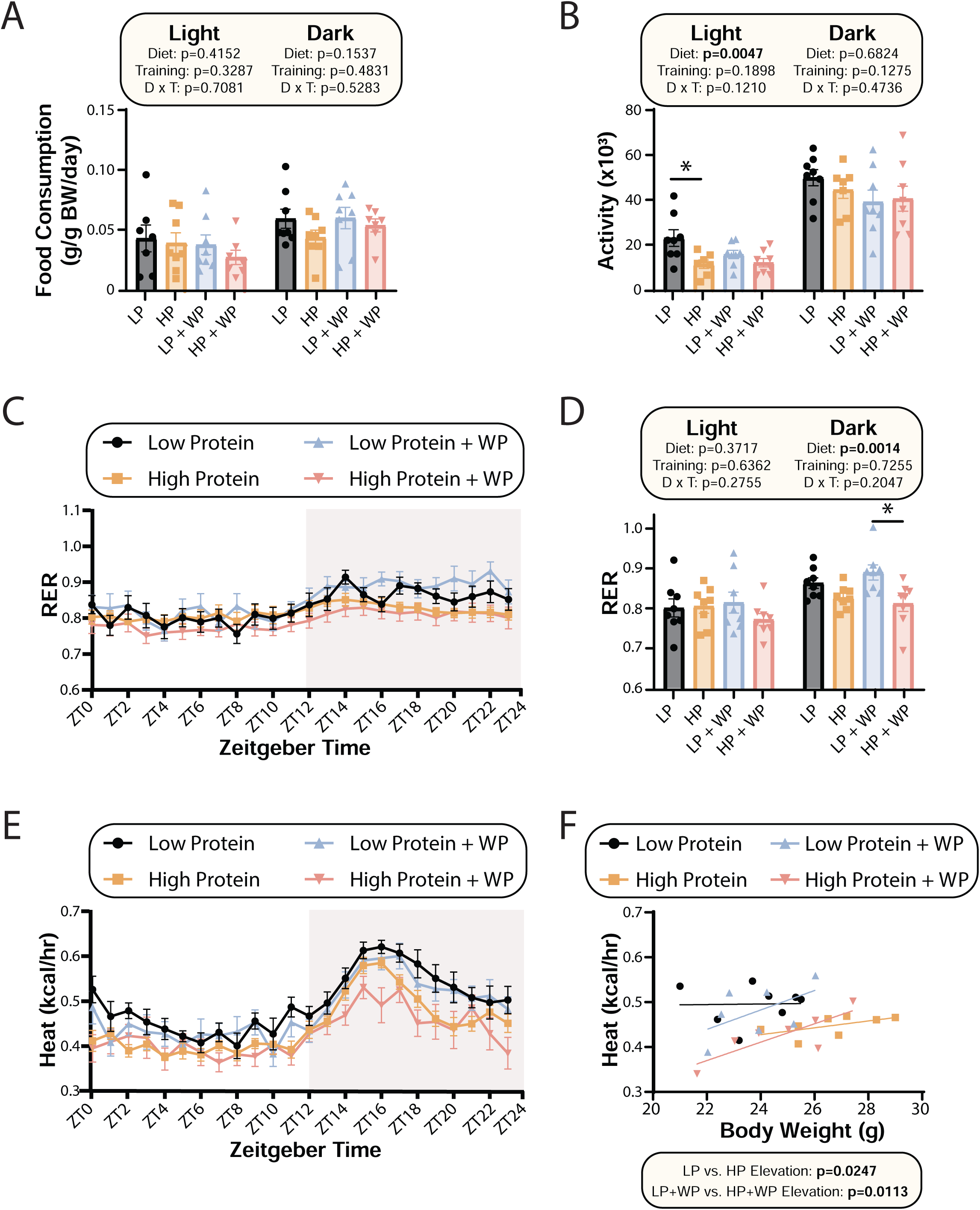
Low protein fed animals have increased energy expenditure regardless of training regimen. (A-F) Metabolic chambers were used to examine metabolic parameters after mice were fed the indicated diets for 8-9 weeks. These included food consumption (A), spontaneous activity (B), Respiratory Exchange Ratio (RER) (C-D), and energy expenditure (E) over a 24 hour period. (F) Energy expenditure (EE) as a function of body weight was calculated (data for each individual mouse are plotted, and slopes and intercepts were calculated using ANCOVA). n=7-8 mice/group. (A, B, D) Statistics for the overall effects of diet, training and the interaction represent the p value from a two-way ANOVA conducted separately for the light and dark cycles; *p<0.05 from a Sidak’s post-test examining the effect of parameters identified as significant in the 2-way ANOVA. Data represented as mean ± SEM.

**Supplemental Figure 2.**
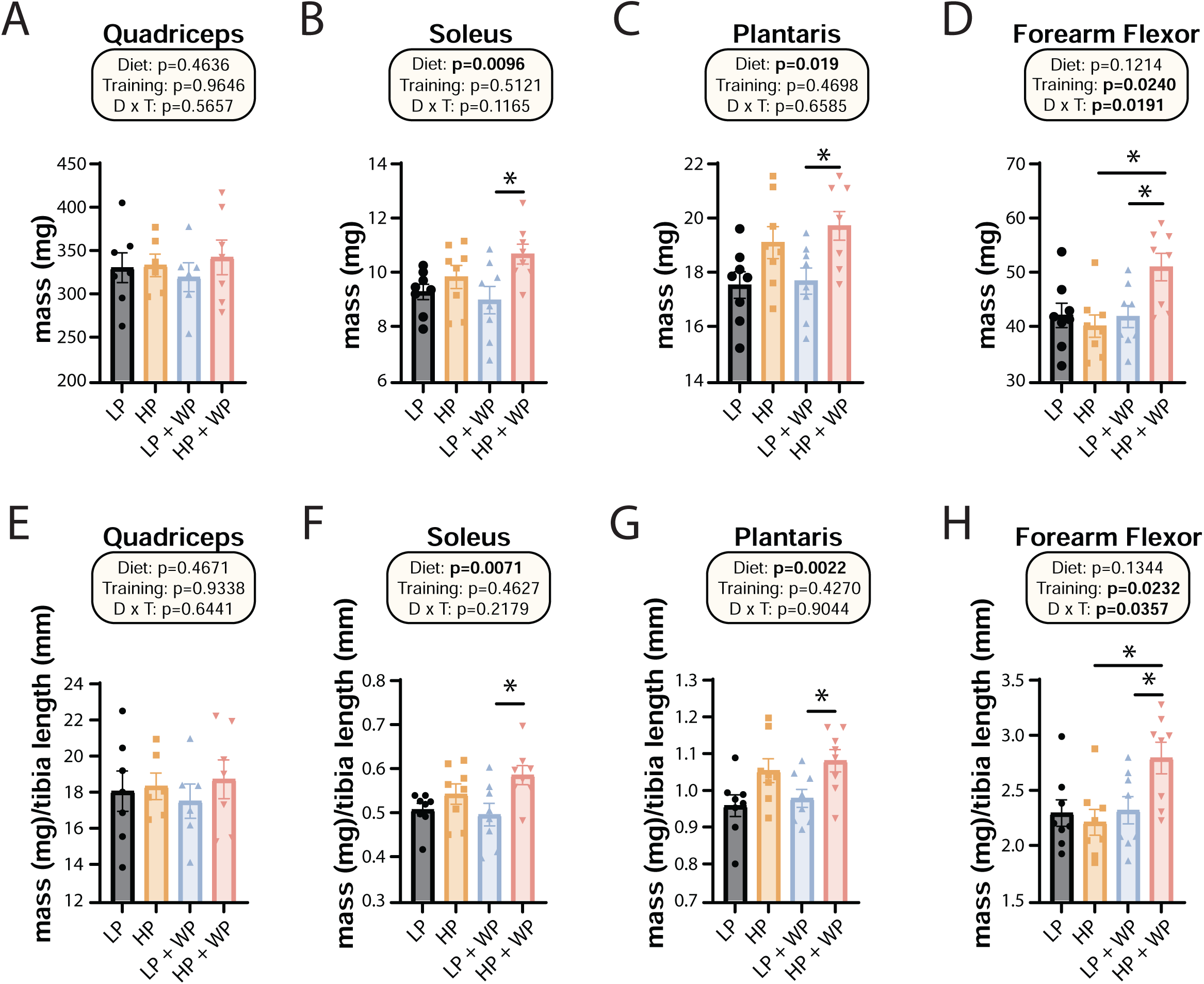
Weight pulling and high protein diet increase the mass of specific muscles. (A-L) The mass of muscles was measured in absolute terms (A-D) and normalized to tibia length (E-H). The muscles measured were the quadriceps (A, E), soleus (B, F), plantaris (C, G) and forearm flexor complex (D, H). n=6-8 mice/group. Statistics for the overall effects of diet, training and the interaction represent the p value from a two-way ANOVA, *p<0.05, Sidak’s post-test examining the effect of parameters identified as significant in the 2-way ANOVA. Data represented as mean ± SEM.

